# Folic acid, but not folate, regulates different stages of neurogenesis in the ventral hippocampus of adult female rats

**DOI:** 10.1101/477448

**Authors:** Wansu Qiu, Aarthi R. Gobinath, Yanhua Wen, Jehannine Austin, Liisa A.M. Galea

## Abstract

Folate is an important regulator of hippocampal neurogenesis, and *in utero* spinal cord development. Both high levels of folic acid and low levels of folate can be harmful to health, as low levels of folate have been linked to several diseases while high folic acid supplements can mask a vitamin B12 deficiency. Depressed patients exhibit folate deficiencies, lower levels of hippocampal neurogenesis, elevated levels of homocysteine, and elevated levels of the stress hormone, cortisol, which may be inter-related. Here, we are interested in whether different doses of natural folate or synthetic folic acid diets can influence neurogenesis in the hippocampus, levels of plasma homocysteine, and serum corticosterone in adult female rats. Adult female Sprague-Dawley rats underwent dietary interventions for 29 days. Animals were randomly assigned to six different dietary groups: folate deficient + succinylsulfathiazole (SST), low 5-methyltetrahydrofolate (5-MTHF), low 5-MTHF + (SST), high 5-MTHF + SST, low folic acid, and high folic acid. SST was added to a subset of the 5-MTHF diets to eliminate folic acid production in the gut. Before and after dietary treatment, blood samples were collected for corticosterone and homocysteine analysis, and brain tissue was collected for neurogenesis analysis. High folic acid and low 5-MTHF without SST increased the number of immature neurons (doublecortin-expressing cells) within the ventral hippocampus compared to folate deficient controls. Low 5-MTHF without SST significantly increased the number of immature neurons compared to low and high 5-MTHF + SST, indicating that SST interfered with elevations in neurogenesis. Low folic acid and high 5-MTHF+SST reduced plasma homocysteine levels compared to controls, but there was no significant effect of diet on serum corticosterone levels. Low folic acid and high 5-MTHF+SST reduced the number of mature new neurons in the ventral hippocampus (BrdU/NeuN-positive cells) compared to folate deficient controls. Overall, folic acid dose-dependently influenced neurogenesis, with low levels decreasing but high levels increasing, neurogenesis in the ventral hippocampus, suggesting this region, which is important for regulating stress, is particularly sensitive to folic acid in diets. Furthermore, the addition of SST negated the effects of 5-MTHF to increase neurogenesis in the ventral hippocampus.

## 1. INTRODUCTION

Folate is one of the natural B-complex vitamins, is responsible for one-carbon metabolism^1^, and converts homocysteine to methionine^2^. This process of reducing homocysteine levels is crucial for DNA synthesis and overall health. The human body is incapable of making folate endogenously. Thus, folate must be obtained through diet, such as by eating leafy greens, legumes, and citrus fruits^3^. Folic acid is a synthetic folate analog and is added to several breads and cereal products to fortify folate levels within the general population. There are however, microorganisms within the human gut microbiome that are capable of synthesizing folic acid from diet^4^. The antibacterial agent, succinylsulfathiazole (SST) is commonly added to folate diets in pre-clinical research studies to prevent folic acid synthesis through the gut microbiome^5^. Thus, SST allows for a controlled dose of folate, without the potentially confounding effects of folic acid, in experimental groups. However, because the addition of SST itself may disrupt neural processes via the gut microbiome. Particularly, germ free with no commensal gut microbiome mice show altered neurogenesis^6^, thus one of the goals of the study was to compare how dietary folate with versus without SST supplementation affected adult female rat neurogenesis.

The world health organization (WHO) recommends the use of folic acid supplements to food products in part to help prevent fetal neural tube development defects during embryonic development^7,8^. However, a number of countries have not complied with the WHO recommendations or have discontinued these supplements^9^. There are differences between natural folate such as 5-methyltetrahydrofolate (L-methylfolate; 5-MTHF) and synthetic folic acid, including the internal chemical structure that leads to different processes of metabolism and different pathways in homocysteine clearance (reviewed in ^10^). For example, folic acid is more bioavailable than natural folate^11^ in diet. In addition, overconsumption of folic acid can lead to excessive levels of unmetabolized folic acid accumulating in blood^12^. Increased and prolonged exposure to unmetabolized folic acid may confer some toxic effects in older female populations^13^. While there are numerous studies supporting folate supplementation in women for the benefit of child development, studies rarely examine whether 5-MTHF or folic acid can affect women’s health. This study aimed to address this gap and directly compare how different doses of both 5-MTHF and folic acid affect endocrine and neural outcomes in adult female rats.

Folate is an important regulator of neuroplasticity, including neurogenesis, during development^14,15^ and during adulthood^16,17^. Particular attention has been paid to folate deficiencies during pregnancy as it is detrimental for fetal nervous system development in utero^18^. In adult mammals, the hippocampus retains the ability to generate new neurons throughout life^19,20^. Although less well studied, folate levels also influence hippocampal neurogenesis in adult or aging populations^16,17^. However, to our knowledge no studies have been conducted examining neurogenesis in adult female rodents after folate manipulations and this study sought to rectify this deficiency in the literature. Metabolites of the 5-MTHF and folic acid metabolic cycle contribute to the formation of purine rings and the conversion of uracil to thymidine for DNA synthesis^21^. Elevated homocysteine levels due to folate deficiencies are associated with cell death and DNA damage^22–24^ that may lead to reduced neurogenesis. In this study we used the endogenous marker, doublecortin (DCX), as it is expressed in immature neurons up until 21-30 days in rats^25^ to examine the effects of 5-MTHF or folic acid in immature neurons. In addition, we examined neurogenesis via the survival of 28-day old bromodeoxyuridine (BrdU)/NeuN cells that were produced and survived under these dietary conditions.

Most studies have examined the effect of folate deficiencies on neurogenesis rather than folate supplementation. Previously, Kronenberg and colleagues^16^ showed that in aged mice (sex not specified), a chronic folic acid-deficient diet for 3 months reduced the number of immature (DCX) neurons in the dentate gyrus (DG) of the hippocampus compared to animals given a folic acid enriched diet. In a separate study, one-month-old (juvenile) male mice maintained on a folate-deficient standard rodent chow for 3.5 months showed reduced adult hippocampal cell proliferation and survival of 18-day old BrdU+ cells (cells that would have been produced and survived in the last 18 days of the diet) compared to a control diet^17^. While these reports show how folate and folic acid deficiencies can reduce neurogenesis and short-term cell survival, no research to our knowledge has specifically investigated the effects of folate dietary interventions on adult hippocampal neurogenesis throughout the diet in healthy female rodents. Furthermore, 5-MTHF diets are thought to have advantages over folic acid diets, including lessening the risk for masking a vitamin B12 deficiency and preventing potential negative effects of unabsorbed folic acid^10^. Here we sought to identify whether 5-MTHF or folic acid at different doses would be more effective in influencing adult hippocampal neurogenesis in females.

Folate and folic acid may have antidepressant properties as assessed in both clinical and preclinical studies^26^. For example, women are more likely to present with perinatal depression when they are folate deficient^27^. In addition, female rodents express less depressive-like behaviour with higher doses of folic acid^28^ and, folic acid can eliminate depressive-like endophenotypes in female mice that had undergone a corticosterone-induced model of depression^29^, which was comparable to pharmacological antidepressant treatment. Depression is associated with reduced hippocampal neurogenesis^30^, elevated cortisol levels^31^, and lower levels of folic acid and elevated homocysteine^32,33^. One mechanism by which folic acid may contribute to mood elevation is through its effects on stress hormones via homocysteine metabolism^34^. For example, acute restraint stress increased homocysteine levels in female rats^35^, and 7 days of folic acid supplementation (30mg/kg) slightly, but not significantly, reduced corticosterone levels in female mice after three weeks corticosterone treatment^29^. These findings suggest that folic acid may elicit antidepressant effects by reducing homocysteine and modulating the corticosterone system. Thus, in the present study we examined homocysteine and corticosterone levels with folate and folic acid diets.

Few studies have studied the differential effects of folic acid versus folate dietary supplementation and thus this study was designed to examine differences between the two diets at different doses on hippocampal neurogenesis, plasma homocysteine and serum corticosterone levels in adult female rats. As folic acid can be synthesized in the gut and not elsewhere in the body, here we used 5-MTHF diets with or without SST to better understand the effects of folate supplementation alone without folic acid. We expect that there would be differential effects of 5-MTHF versus folic acid to increase neurogenesis in the hippocampus of adult female rats that may be associated with changes in plasma homocysteine and corticosterone. Due to the lack of data on the effects of the addition of SST in the presence of folate diets, and how it affects neurogenesis, here we will compare how low folate diets with or with SST will influence neurogenesis, plasma homocysteine and corticosterone, to determine if at a lower dose of folate SST can negate the effects of dietary folate.

## 2. MATERIALS AND METHODS

### 2.1 Animals

Thirty adult female Sprague-Dawley (Charles River, Quebec) rats (7 months old) were individually housed in transparent polyurethane bines (24 x 16 x 46 cm) with aspen chip bedding, in order to ensure proper monitoring of food consumption. Rats were maintained in a 12 h: 12 h light/dark cycle (lights on at 07:00) and provided rat chow and tap water *ad libitum*. All protocols were in accordance with ethical guidelines set by Canada Council for Animal Care and were approved by the University of British Columbia Animal Care Committee. For an overview of experimental procedures, refer to Figure 1.

**Figure 1.**
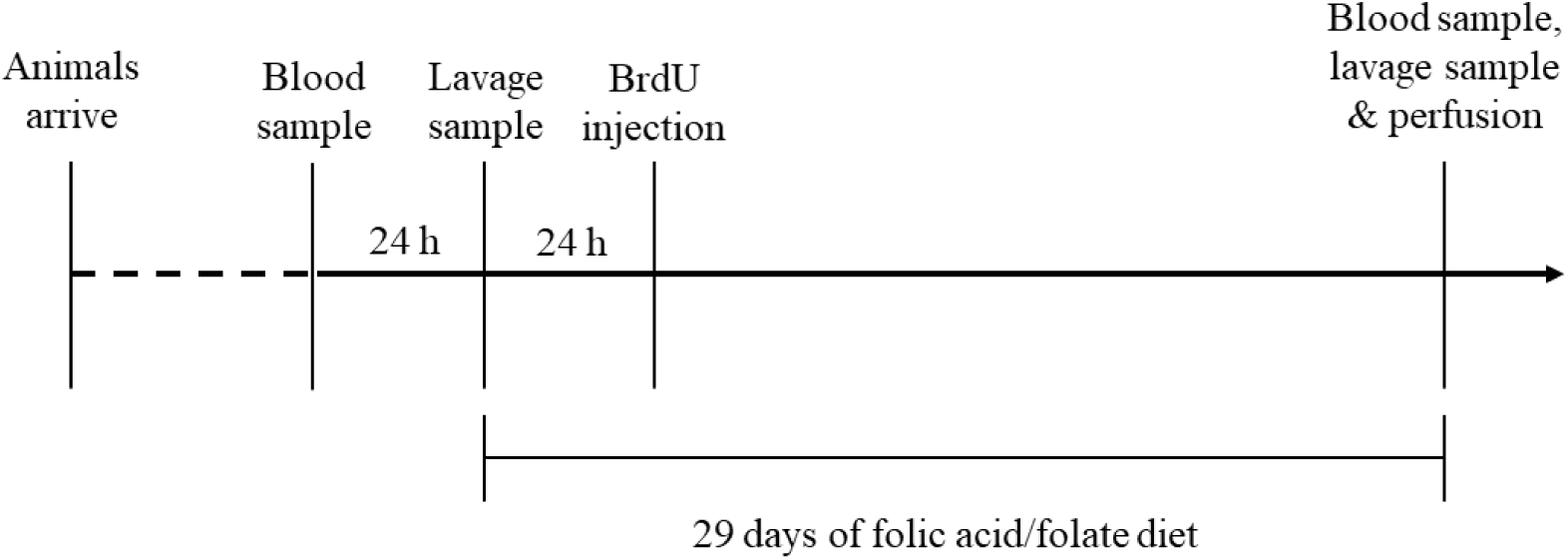
Timeline of Experiment.

### 2.2 Dietary Interventions

Animals were randomly assigned to one of the following diets (n=5/group; total n=30): 1. Folate deficient + 1% succinylsulfathiazole (SST; FD+) that will serve as the control group, 2. Low 5-methyltetrahydrofolate (5-MTHF) + SST was the low folate + SST group (L5-MTHF+), 3. Low 5-MTHF without SST was the low folate group (L5-MTHF), 4. High 5-MTHF + 1% SST was the high folate + SST group (H5-MTHF+), 5. Low folic acid (LFA), 6. High folic acid (HFA). Specifically, 5-MTHF was purchased from Vitacost.com, Boca Raton, FL, USA and formulated with a control folate and folic acid deficient diet (TD.06691) by Harlan Laboratories, Inc., Madison WI, US, to create the 5-MTHF diets. Folic acid diets were formulated directly by Harlan Laboratories, Inc., Madison WI, US with the control diet (TD.06691). The antibacterial agent, SST was added to folate diets and the control diet to prevent folic acid synthesis through the gut microbiome^5^ by Harlan laboratories Inc., Madison WI, US, directly. In animals with SST treatment, it was assumed that there would not be any *de novo* folic acid synthesis by gut bacteria, as it is commonly used to induce a folic acid deficient diet for rodents^36^. The control diet does not contain any folate or folic acid, all other diets were identical with the exception of the additional doses of 5-MTHF or folic acid per treatment group. All other vitamins as well as caloric value remained the same among all 6 diets. For an overview of details regarding the diets, refer to Table 1. Diets were given for 29 days. Doses were chosen based on studies in which similar doses showed efficacy in increasing neurogenesis in male rats in a model of cerebral ischemia^37^. Food consumption was monitored (food weighed) for weekly intake of folic acid or folate diets. Body mass was also monitored every 7 d.

**Table 1.**
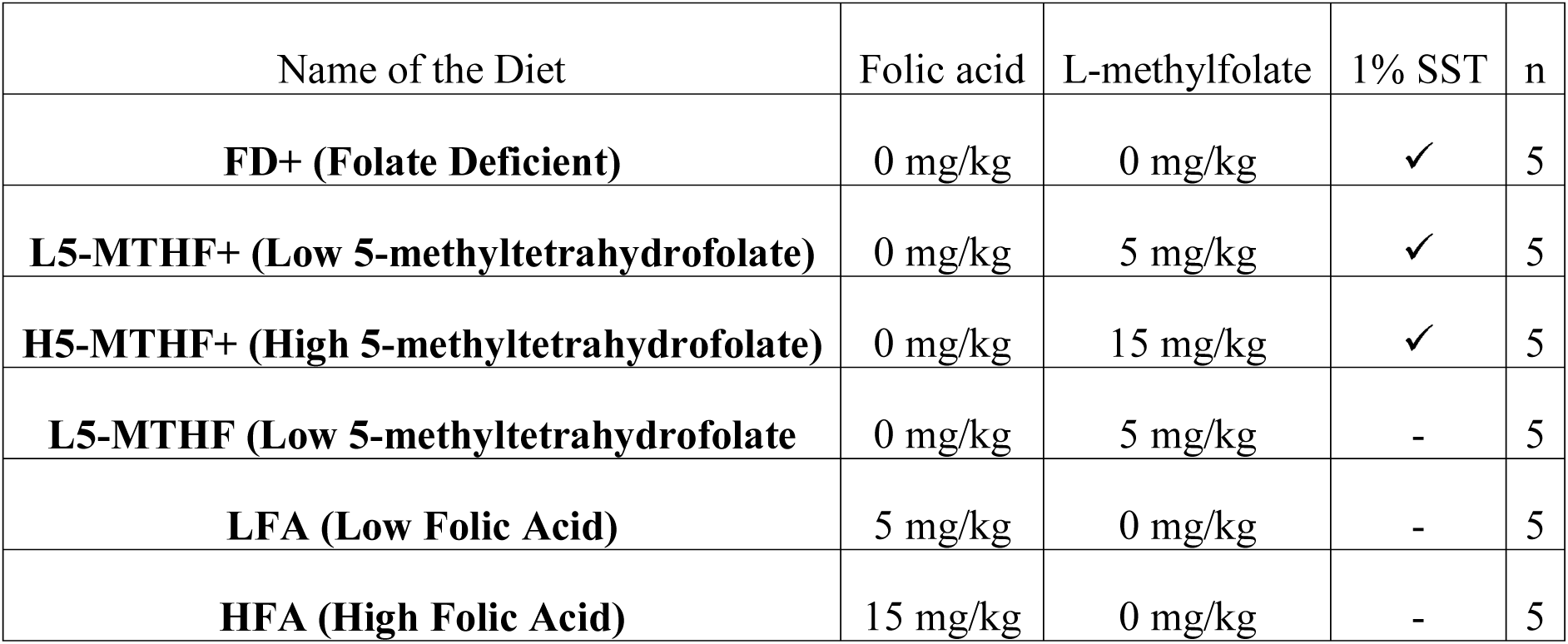
Description of dietary treatments.

| Name of the Diet | Folic acid | L-methylfolate | 1% SST | n |
| --- | --- | --- | --- | --- |
| **FD+ (Folate Deficient)** | 0 mg/kg | 0 mg/kg | ✓ | 5 |
| **L5-MTHF+ (Low 5-methyltetrahydrofolate)** | 0 mg/kg | 5 mg/kg | ✓ | 5 |
| **H5-MTHF+ (High 5-methyltetrahydrofolate)** | 0 mg/kg | 15 mg/kg | ✓ | 5 |
| **L5-MTHF (Low 5-methyltetrahydrofolate)** | 0 mg/kg | 5 mg/kg | - | 5 |
| **LFA (Low Folic Acid)** | 5 mg/kg | 0 mg/kg | - | 5 |
| **HFA (High Folic Acid)** | 15 mg/kg | 0 mg/kg | - | 5 |

### 2.3 Estrous cycle sample and cytology

Vaginal lavage samples were taken on the first day of the diet, and again on the last day of the diet. To determine estrous cycle stages, lavage slides were stained with cresyl violet stain and classified via microscopy by cell phenotypes^38^.

### 2.4 Bromodeoxyuridine (BrdU) preparation

Approximately 24 h after animals began their new diet, animals received a single injection of bromodeoxyuridine (BrdU; 200 mg/kg dose; i.p.), dissolved in 0.9 saline for a stock solution concentration of 20mg/ml. BrdU is a thymidine analogue and a DNA synthesis marker, which labels dividing cells and their progeny. Thus, newly dividing cells were born in different folate/folic acid environments, allowing us to investigate whether cells born into different folate/folic acid environments would affect subsequent survival of these BrdU-positive (BrdU+) cells. Animals were euthanized 28 days later, therefore BrdU tagged cells were 28 days old.

### 2.5 Blood collection, corticosterone, and homocysteine analysis

Twenty-four hours prior to dietary interventions and again after 29 days of dietary intervention, blood samples were collected via tail vein. All blood samples were collected within 3 min of touching the cage.

Plasma samples were collected with the anticoagulant EDTA (37.5 mg/ml) dissolved in deionized water. Plasma blood was then centrifuged at 10,000 g for 15 min immediately. Serum blood samples were stored overnight at 4ºC to allow blood to clot completely, and then centrifuged at 10,000 g for 15 min. The serum and plasma were collected and stored at −20ºC until radioimmunoassay.

Total CORT (bound and free) was measured on the serum samples using the ImmuChem Double Antibody 125I radioimmunoassay Kit (MP Biomedicals, Solon, OH, USA). The antiserum cross-reacts 100% with CORT, 0.34% with deoxycorticosterone, 0.05% with cortisol, and does not cross-react with dexamethasone (<0.01%). All reagents were halved, and samples run in duplicates. Percent change in serum CORT was calculated following the formula ((CORT levels after treatment – CORT levels prior to treatment)/CORT levels prior to treatment) * 100.

Plasma homocysteine levels were measured by liquid chromatography-tandem mass spectrometry in the Analytical Core for Metabolomics and Nutrition laboratories located at BC Children’s Hospital Research Institute at the University of British Columbia as described by Dominguez-Salas et al.^39^ once after dietary treatment.

### 2.6 Tissue Collection

On Day 29, rats were then weighed and given an overdose of Sodium Pentobarbitol (Euthanyl). Adrenal glands and ovarian tissue were collected and weighted. Rats were then transcardially perfused with 60 ml cold saline followed by 120 ml cold 4% paraformaldehyde (in 0.1M phosphate buffer). Brains were then extracted and post-fixed containing 4% paraformaldehyde overnight at 4ºC. Brains were then transferred to 30% sucrose in phosphate buffer at 4ºC. Brains were rapidly frozen and sectioned using a freezing microtome (Leica, Richmond Hill, ON, Canada) at 30 µm in a series of 10. Sections were stored in 30 % ethylene glycol/ 20% glycerol in phosphate buffer; Sigma) and stored at −20ºC until processing.

### 2.7 Immunohistochemistry

#### Doublecortin (DCX)

DCX is expressed in immature neurons^40^ between 4 hours to 21 days after production in rats^25^ with longer timelines in mice^41^. In our case, DCX-expressing cells would have been exposed to dietary manipulations. The morphology of DCX-expressing cells has been used to show the maturity stages of these new neurons^42^ (proliferative, intermediate, and post-mitotic) and generally the more mature morphology will indicate a longer duration of exposure of the DCX-expressing cells to the diet. Thus, here we provide the number of DCX-expressing cells and also the morphology of those cells to determine whether morphological phenotype different with dietary condition. Immunohistochemistry for DCX was conducted as previously described ^34,35^. Briefly, sections were rinsed 5 x 10 min in 0.1 M phosphate buffered saline (PBS), treated with 0.3% hydrogen peroxide in dH_2_O for 30 min, and incubated at 4 ºC in primary antibody solution: 1:1000, goat anti-DCX (Santa Cruz Biotechnology, Santa Cruz, CA, USA) with 0.04% Triton-X in PBS and 3% normal rabbit serum for 24 h. Sections were then rinsed 5 x 10 min in 0.1 M PBS and transferred to a secondary antibody solution with 1:500, rabbit anti-goat (Vector Laboratories, Burlington, ON, Canada) in 0.1 M PBS for 24 h at 4ºC. Then, sections were washed 5 x 10 min in 0.1 M PBS and incubated in ABC complex (ABC Elite Kit; 1:1000; Vector) for 4 h. Sections were then washed in 0.175 M sodium acetate buffer 2 x 2 min. Finally, sections were developed using diaminobenzidine in the presence of nickel (DAB Peroxidase Substrate Kit, Vector), mounted on slides, and dried. Sections were then counterstained with cresyl violet, dehydrated, and cover-slipped with Permount (Fisher Scientific, Hampton, NH, USA).

#### Bromodeoxyuridine (BrdU)

To examine the influence of dietary interventions on cytogenesis and complement our DCX data with the DNA synthesis marker, BrdU. BrdU can be used as a marker of cell proliferation or survival of new cells depending on the timeline between injection and perfusion^43^. Here, we are examining the influence of diet on cell survival as we injected BrdU on day 1, 24 h after dietary supplementation and perfused the animals 28 days later. Sections were rinsed 3 x 10 min in 0.1 M Tris-buffered saline (TBS), treated with 0.6% hydrogen peroxide in dH_2_O for 30 min, and washed 3 x 10 min in 0.1 M TBS. Sections were then incubated in 2N hydrochloric acid at 37 ºC for 30 min. And then incubated in 0.1M borate buffer for 10 min. Sections were then rinsed 3 x 10 min in 0.1 M TBS and blocked with 0.3% Triton-X and 3% normal horse serum (MilliporeSigma, MA, USA) mixed in 0.1 M TBS (TBS+). Sections were then incubated at 4 ºC in primary antibody solution: 1:200, mouse anti-BrdU (Roche, Basel, Switzerland) mixed in TBS+ solution for 30-48 h. Sections were then rinsed 3 x 10 min in 0.1 M TBS and transferred to a secondary antibody solution with 1:200 anti-mouse IgG (Vector Laboratories, Burlington, ON, Canada) in TBS+ for 4 h at 4ºC. Then, sections were washed 3 x 10 min in 0.1 M TBS and incubated in ABC complex (ABC Elite Kit; 1:1000; Vector) for 1.5 h. Sections were then washed 3 x 10 min in 0.1 M TBS. Finally, sections were developed using diaminobenzidine in the presence of nickel (DAB Peroxidase Substrate Kit, Vector), washed 3 x 10 min in 0.1 M TBS, mounted on slides, and dried. Sections were then counterstained with cresyl violet, dehydrated, and cover-slipped with Permount (Fisher Scientific, Hampton, NH, USA).

#### BrdU/NeuN double labelling

BrdU/NeuN immunofluorescent double labelling was used to quantify the number of new cells (BrdU+) that have differentiated into neurons (NeuN) and have survived within the hippocampus to examine neurogenesis (Figure 4E)^44^. All brain slices were rinsed in tris-buffered saline (TBS) 3 x 10 min and then blocked with TBS + 0.3% Triton-X and 3% normal donkey serum (MilliporeSigma, MA, USA; TBS+) for 30min prior to all primary and secondary antibody incubations. All primary and secondary antibody solutions were made with TBS + 0.1% Triton-X and 3% NDS. Briefly, tissue slices were incubated in primary 1:500 mouse anti-NeuN (MilliporeSigma, MA, USA) for approximately 48hrs at 4°C. After, tissue slices were incubated in secondary donkey anti-mouse ALEXA 488 (Invitrogen, Burlington, ON, Canada) overnight at 4°C. Brain slices were then rinsed 3 x 10 min in TBS, fixed in 4% paraformaldehyde for 10 min, and then rinsed 2 x 10 min in 0.9% saline solution. Brain sections then were incubated in 2N hydrochloric acid for 30min at 37°C and incubated in primary rat anti-BrdU (Abchem Inc, Dorval, QC, Canada) for approximately 48hrs at 4°C. After, sections were incubated in secondary donkey anti-rat ALEXA 594 (Invitrogen, Burlington, ON, Canada) overnight at 4°C. Finally, sections were rinsed in TBS 3 x 10 min prior to being mounted onto microscope slides and cover-slipped with PVA DABCO.

### 2.8 Microscopy

#### Doublecortin (DCX)

Hippocampal slices (1/10^th^) were exhaustively counted for DCX-expressing cells using the 40x objective of an Olympus C×22LED brightfield microscope. Cells were counted separately in the dorsal region (−2.76 mm to −4.68mm below bregma) and in the ventral region (−5.52 mm to −6.60 mm below bregma) as previous studies have shown that these areas can serve different functions^45^. For exhaustive DCX-expressing cell counts, all sections were counted per dorsal (∼8-11 sections) and ventral region (∼7-14 sections; no difference in section numbers between dietary groups), and then total counts per region were recorded. The total cell count was then multiplied by 10 (number of slices) to obtain an estimate of the total number of DCX-expressing cells in the region^46^. Areas of the dentate gyrus were quantified from digitized images using ImageJ (NIH, Bethesda, MD, USA), and volumes were calculated using Cavalieri’s principle^47^. Sixty cells (n = 30 within the dorsal regions; n = 30 within the ventral region) were randomly selected and were classified by type into 1 out of 3 developmental stages of DCX-expressing neurons (proliferative, intermediate, and postmitotic) based on morphological attributes for cell maturity analysis^48^. The percentage of these DCX cells in each morphological state were recorded. Experimenters were blind to the treatment conditions when conducting microscopy analyses.

#### Bromodeoxyuridine (BrdU)

BrdU+ cells were quantified in all dorsal sections (−2.76 mm to −4.68mm below bregma) and all ventral sections (−5.52 mm to −6.60 mm below bregma) using the 100x immersion objective of a Nikon E600 light microscope in every 10^th^ section of the granule cell layer including the subgranular zone (∼5μl of cells between the granule cell layer and hilus). For exhaustive BrdU+ cell counts, all sections were counted per dorsal (∼5-11 sections) and ventral region (∼6-12 sections; no difference in section numbers between dietary groups). We used a modified optical dissector method^49,50^ to estimate the total number of BrdU+ and DCX-expressing cells, as has been used before^51–55^. Total BrdU+ cell counts were determined by multiplying by 10.

#### BrdU/NeuN

Fifty BrdU+ cells were randomly selected between dorsal (n = 25 within the dorsal region) and ventral sections (n = 25 within the ventral region) using the 60x immersion objective of an Olympus FV1000 confocal microscope. The percentage of BrdU+ cells that also expressed NeuN were quantified. A neurogenesis index factor was calculated by multiplying the percentage of BrdU+ cells that also expressed NeuN with the total raw count number of BrdU+ cells counted as has been done in previous studies^41,56^.

### 2.9 Data Analyses

All data were analyzed using one-way analysis of variance (ANOVA) unless otherwise specified with diet (FD+, L5-MTHF+, L5-MTHF, H-5MTHF+, LFA, and HFA) as between-subject factors. Adrenal mass, and ovarian mass were analyzed using one-way analysis of covariance (ANCOVA) using body mass as a covariate. DCX-expressing cells and relative dentate gyrus volume, BrdU+ cells or BrdU/NeuN-positive (BrdU/NeuN+) cells were analyzed using repeated measures ANOVA with diet as between-subject factors, and region (dorsal, ventral) as the within-subjects factor. An additional within-subject factor of developmental stages (proliferative, intermediate, and post-mitotic) for cell maturity analysis for DCX-expressing cells. Effect sizes are reported for significant effects. Post hoc comparisons used Fisher’s least significant difference (LSD) test. *A priori* comparisons were subjected to Bonferroni correction. Pearson product-moment correlations were conducted between body mass, relative organ mass, serum CORT, dorsal and ventral DCX-expressing cells, dorsal and ventral BrdU+ cells, and homocysteine levels. All data were analyzed using Statistica software (v. 9, StatSoft, Inc., Tulsa, OK, USA). All effects were considered statistically significant if *p* ≤ 0.05, trends are discussed if *p* ≤ 0.10. Outliers were eliminated if higher or lower than 2 standard deviations from the mean. This happened twice, one case outlier was excluded from the FD+ group when analyzing serum CORT levels and another outlier was excluded from the L-5MTHF+ group when analyzing BrdU/NeuN cells.

## 3. RESULTS

### 3.1 High folic acid and folate increased adrenal mass, and low folic acid and high folate diets reduced homocysteine levels diets but did not influence body mass, ovarian mass, or percent change in serum CORT levels

The HFA group had a significant increase in adrenal mass when accounting for the covariate body mass compared to FD+ (*p* = 0.003, Cohen’s *d* = 2.900) and trended towards significance in higher adrenal mass compared to the LFA group (*p* = 0.086). H-5MTHF+ group had a significant increase in adrenal mass compared to FD+ (*p* = 0.036, Cohen’s *d* = 1.575) and L5-MTHF+ (*p* = 0.044, Cohen’s *d* = 1.655; main effect of diet: F(5, 23) = 3.185, *p* = 0.025, □_p_^2^ = 0.409), but there were no other significant effects on adrenal mass (all *p’s* ≥ 0.159; Figure 2A). Diet did not statistically influence body mass or ovarian mass (all *p*’s ≥ 0.480). Diet did not statistically influence percent change in serum CORT levels (*p* = 0.169). Diet did not statistically influence the relative 1/10^th^ volume of the hippocampus (*p* = 0.811).

**Figure 2.**
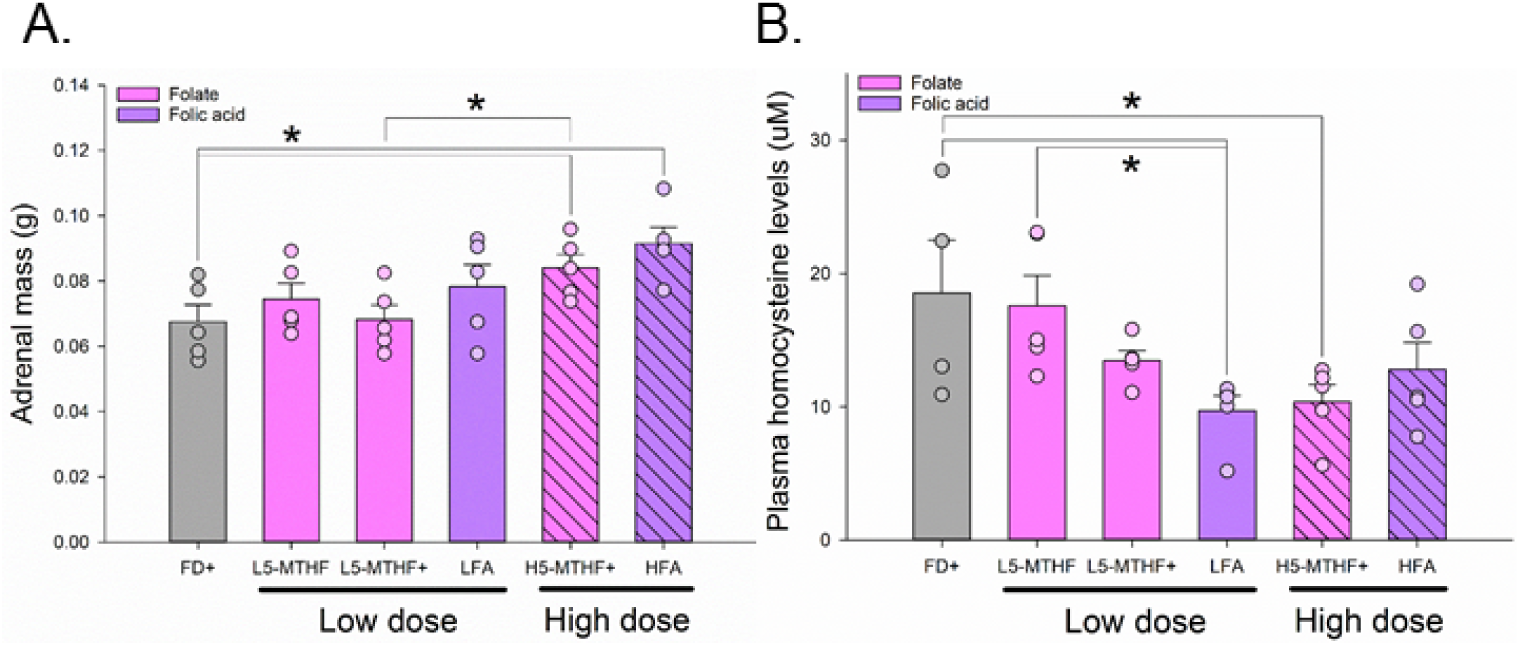
**A.** Mean and + standard error of the mean (SEM) of adrenal mass, and individual data points of adrenal mass dependent on dietary interventions. High folic acid (HFA) and high folate + succinylsulfathiazole (H5-MTHF+) animals showed increased adrenal mass when controlling for final body mass compared to folate deficient (FD+) animals. H5-MTHF+ animals also had larger adrenal mass compared to low folate + succinylsulfathiazole (L5-MTHF+), * indicate significance at *p* = 0.05. **B.** Mean and + SEM of plasma homocysteine levels (uM) and individual data points of homocysteine levels. Low folic acid (LFA) animals showed a significant decrease in homocysteine levels compared to FD+ controls and low folate (L5-MTHF) animals. H5-MTHF+ animals also showed a significant decrease in homocysteine levels compared to FD+ animals, * indicate significance at *p* = 0.05

Animals did not differ significantly in estrous cycle stages at the beginning or at the end of the experiment (*p* = 0.313, *p* = 0.637, respectively) nor was estrous cycle stage affected by dietary treatment (*p* = 0.493). Overall, estrous cycle stage did not influence the number of DCX-expressing or BrdU+ cells (*p* = 0.254, *p* = 0.769, respectively).

LFA animals showed a significantly lower homocysteine levels at the end of dietary treatment compared to FD+ controls (*p* = 0.006, Cohen’s *d* = 1.501; main effect of diet: F(5, 23) = 3.177, *p* = 0.025, □_p_^2^ = 0.408; Figure 2B) and L5-MTHF animals (*p* = 0.009, Cohen’*d* = 1.369). H5-MTHF+ animals showed a significant reduction in homocysteine levels compared to FD+ (*p* = 0.011, Cohen’s *d* = 1.956). HFA animals were trending towards significance in a reduction in homocysteine levels compared to FD+ animals (*p* = 0.064), no other significant comparisons were found (all *p*’s ≥ 0.100).

### 3.2 High folic acid and low folate (without SST) increased DCX-expressing cells in the ventral dentate gyrus

The HFA diet significantly increased the number of ventral hippocampal DCX-expressing cells compared to the FD+ (*p* = 0.008, Cohen’s *d* = 0.894), LFA (*p* = 0.002, Cohen’s *d* = 1.094), and H5-MTHF+ (*p* = 0.014, Cohen’s *d* = 0.909) diets. L5-MTHF diet significantly increased the number of ventral hippocampal DCX-expressing cells compared to FD+ (*p* = 0.012, Cohen’s *d* = 0.719), L5-MTHF+ (*p* < 0.001, Cohen’s *d* = 1.164), and LFA (*p* = 0.006, Cohen’s *d* = 0.888) diets (Figure 3A). HFA diet significantly increased the number of dorsal hippocampal DCX-expressing cells compared to H5-MTHF+ diet (*p* = 0.032, Cohen’s *d* = 0.770; Figure 3B. A significant interaction between diet and dentate gyrus region, F(5, 24) = 2.652, *p* = 0.048, □_p_^2^ = 0.356, and a significant main effect of dentate gyrus region, F(1, 24) = 16.701, *p* < 0.001, □_p_^2^ = 0.410. No other significant comparisons were found (all *p*’s ≥ 0.102).

**Figure 3.**
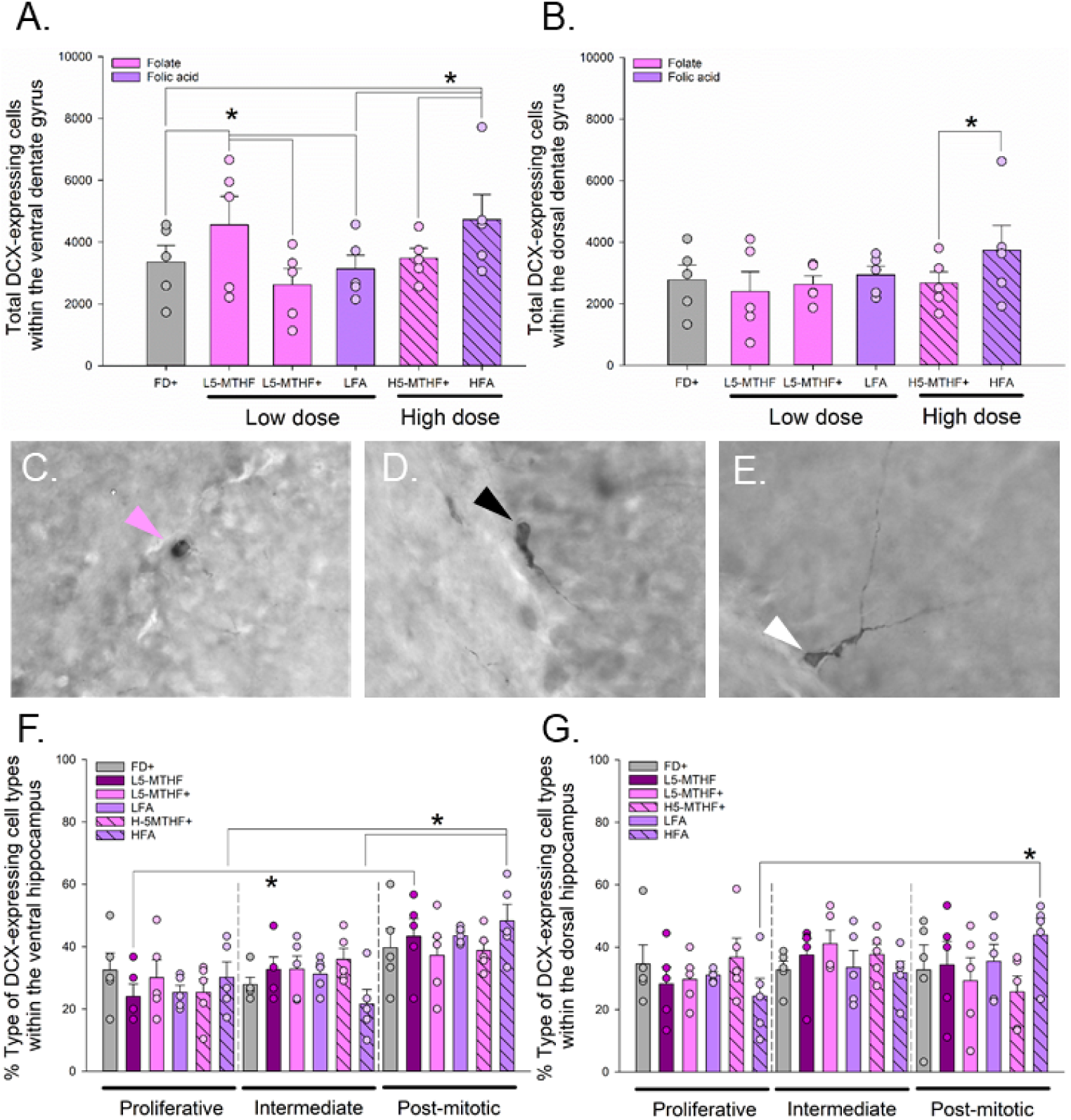
**A.** Mean and + standard error of the mean (SEM) of total doublecortin (DCX)-expressing cell counts, and individual data points of DCX-expressing cell counts dependent on dietary interventions within the ventral dentate gyrus region of the hippocampus. High folic acid (HFA) group showed significantly more DCX-expressing immature neurons within the ventral region compared to folate deficient (FD+) controls, low folic acid (LFA) and high folate + succinylsulfathiazole (H5-MTHF+) groups. Low folate without SST (L5-MTHF) group showed higher DCX-expressing immature neurons compared to the FD+ controls, low folate + SST (L5-MTHF+), and LFA group, * indicate significance at *p* = 0.05. **B.** Mean and + SEM of DCX-expressing cell counts, and individual data points of DCX-expressing cell counts dependent on dietary interventions within the dorsal dentate gyrus region of the hippocampus. HFA group showed significantly higher DCX-expressing cell counts compared only to the H5-MTHF+ group, * indicate significance at *p* = 0.05. **C-E.** DCX-expressing cells along the granule cell layer of the hippocampal dentate gyrus, pictures taken at 60x objective magnification. Arrow head indicated DCX-expressing cells (pink: proliferative DCX-expressing cells; black: intermediate DCX-expressing cells; white: post-mitotic DCX-expressing cells). **F.** Mean and + SEM of % type of DCX-expressing cells, and individual data points of % type of DCX-expressing cells within the ventral region of the hippocampus. HFA animals showed significantly more post-mitotic cells compared to proliferative and intermediate cells. L5-MTHF animals showed significantly more post-mitotic cells compared to proliferative cells, * indicate significance at *p* = 0.05. **G.** Mean and + SEM of % type of DCX-expressing cells, and individual data points of % type of DCX-expressing cells within the dorsal region of the hippocampus. HFA animals showed significantly more post-mitotic cells compared to proliferative cells, * indicate significance at *p* = 0.05.

### 3.3 Animals showed significantly higher numbers of post-mitotic doublecortin cells overall, particularly when treated with high folic acid and low folate without SST

Within the ventral dentate gyrus, post hoc analysis indicated a significantly higher percentage of post-mitotic cells compared to both proliferative and intermediate cells with a significant main effect of type of cells, F(2, 48) = 3.701, *p* = 0.032, □_p_^2^ = 0.134, and a significant interaction between type of cells and dentate gyrus region, F(2, 48) = 10.243, *p* < 0.001, □_p_^2^ = 0.299 (Table 2). Within the dorsal dentate gyrus, post hoc analysis indicated a significantly higher percentage of intermediate cells compared to proliferative cells (Table 2). *A priori* post-hoc analysis indicated within the HFA diet group, there were significantly more post-mitotic DCX-expressing cells compared to proliferative cells (*p* = 0.002, Cohen’s d = 1.585) and intermediate cells (*p* < 0.001, Cohen’s d = 2.404) within the ventral dentate gyrus (Figure 3F), and more post-mitotic DCX-expressing cells compared to proliferative cells (*p* = 0.001, Cohen’s d = 1.592) within the dorsal dentate gyrus (Figure 3G). L5-MTHF animals also showed a significantly higher percentage post-mitotic cells compared to proliferative cells only within the ventral region (*p* = 0.001, Cohen’s d = 1.761).

**Table 2.** Description of *p*-values and effect sizes for post-hoc comparisons between dentate gyrus region and type of DCX-expressing cell (proliferative, intermediate, post-mitotic), * indicate significance at *p* = 0.05.

|  |  |  |  | Intermediate | Post-mitotic |
| --- | --- | --- | --- | --- | --- |
| Region | Type of DCX cells | Mean | SD | Dorsal region |  |
| Dorsal | Proliferative | 30.759 | 11.007 | $p = 0.034^*$<br>(Cohen's $d = 0.486$) | $p = 0.225$ |
| | Intermediate | 35.696 | 9.241 | **X** | $p = 0.347$ |
|  | Post-mitotic | 33.544 | 14.549 | **X** | **X** |
|  |  |  |  | Ventral region |  |
| Ventral | Proliferative | 27.887 | 9.834 | $p = 0.287$ | $p < 0.001^*$<br>(Cohen's $d = 1.360$) |
| | Intermediate | 30.3260 | 8.715 | **X** | $p < 0.001^*$<br>(Cohen's $d = 1.182$) |
|  | Post-mitotic | 41.786 | 10.591 | **X** | **X** |

### 3.4 Low folic acid and high folate with SST decreased the number of BrdU/NeuN+ cells within the ventral dentate gyrus

LFA animals showed a significant reduction in ventral BrdU+ cells compared to FD+ (*p* = 0.0161, Cohen’s *d* = 1.533), L5-MTHF (*p* = 0.008, Cohen’s d = 1.286), and HFA (*p* = 0.001, Cohen’s *d* = 2.668) animals, and a trend towards significance in lower BrdU+ cells compared to L5-MTHF+ (*p* = 0.064; Figure 4C). H5-MTHF+ animals showed a significant reduction in BrdU+ cells compared to FD+ (*p* = 0.048, Cohen’s *d* = 1.048) and HFA (*p* = 0.005, Cohen’s *d* = 1.805) animals. In the dorsal hippocampus, LFA animals showed a significant reduction in BrdU+ cells compared to FD+ (*p* = 0.015, Cohen’s *d* = 0.992), L5-MTHF+ (*p* = 0.004, Cohen’s *d* = 1.755), and HFA (*p* < 0.001, Cohen’s *d* = 1.812) animals (Figure 4D). H5-MTHF+ animals showed a significant reduction compared to HFA animals (*p* = 0.019, Cohen’s *d* = 0.992), with a main effect of diet, F(5, 24) = 3.105, *p* = 0.027, □_p_^2^ = 0.393, and main effect of region, F(1, 24) = 16.864, *p* < 0.001, □_p_^2^ = 0.413, but no significant interaction effect between region and diet (*p* = 0.466). No other pair-wise post hoc comparisons (all *p*’s ≥ 0.135).

**Figure 4.**
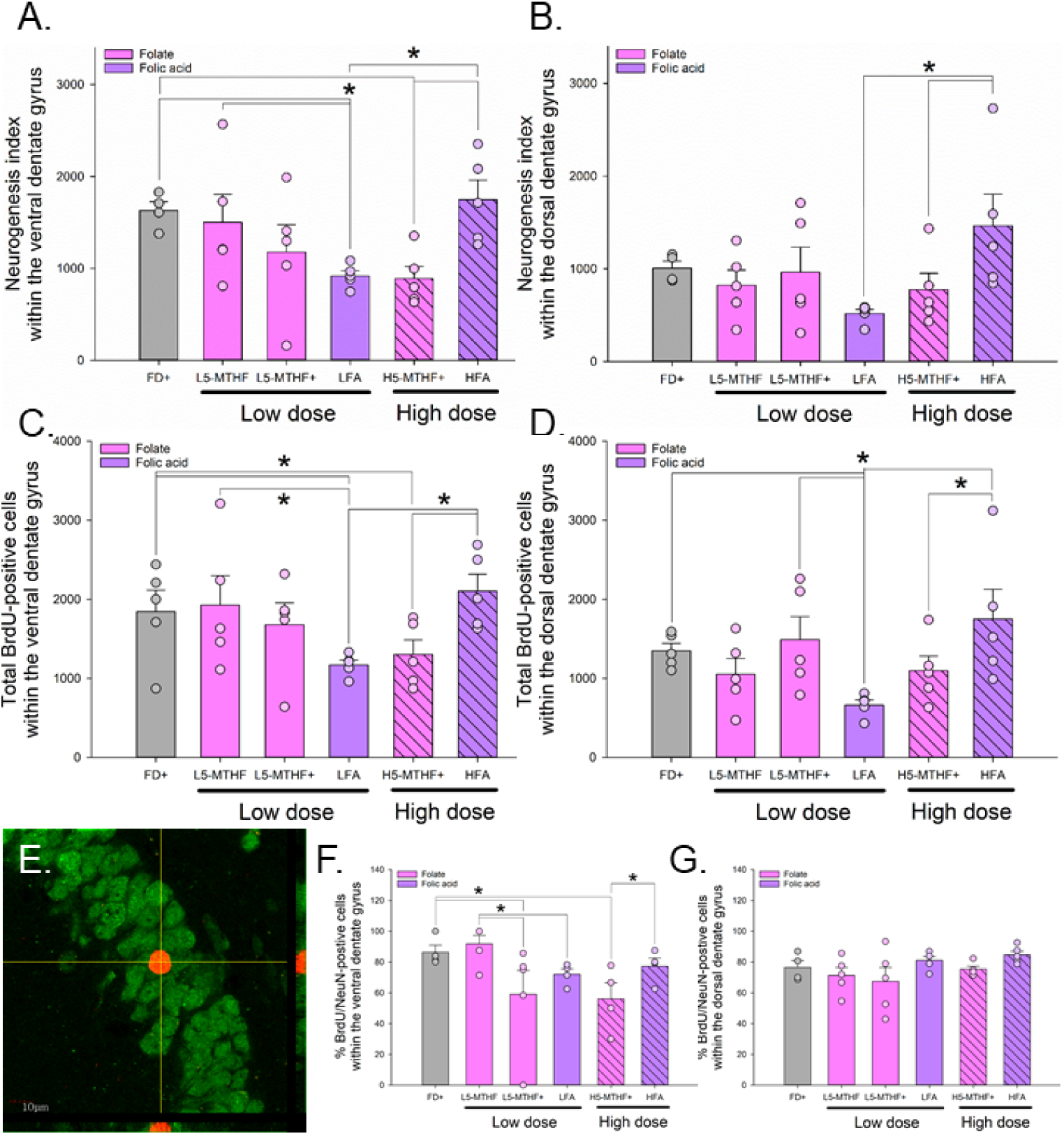
**A.** Mean and + standard error of the mean (SEM) of neurogenesis index factor (percentage of bromodeoxyuridine (BrdU)/NeuN-positive (BrdU/NeuN+) cell counts × total BrdU-positive (BrdU+) cell counts), and individual data points of neurogenesis index factor dependent on dietary intervention within the ventral dentate gyrus region of the hippocampus. High folate + succinylsulfathiazole (H5-MTHF+) decreased neurogenesis compared to folate deficient (FD+) controls, and high folic acid (HFA) animals. Low folic acid (LFA) decreased neurogenesis compared to FD+, low folate (L5-MTHF), and HFA animals, * indicate significance at *p* = 0.05. **B.** Mean and + SEM of neurogenesis index factor, and individual data points of neurogenesis index factor dependent on dietary intervention within the dorsal dentate gyrus region of the hippocampus. HFA animals showed higher neurogenesis compared to LFA and H5-MTHF+ animals, * indicate significance at *p* = 0.05. **C.** Mean and + SEM of total BrdU+ cell counts, and individual data points of BrdU+ cell counts dependent on dietary intervention within the ventral dentate gyrus region of the hippocampus. HFA group showed significantly more BrdU+ cells within the ventral region compared to LFA and H5-MTHF+ groups. LFA group showed significantly less BrdU= cells compared to the L5-MTHF and FD+ groups. H5-MTHF+ showed a significantly lower number of BrdU+ cells compared to FD+ group, *indicate significance at *p* = 0.05. **D.** Mean and + SEM of total BrdU+ cell counts, and individual data points of BrdU+ cell counts dependent on dietary interventions within the dorsal dentate gyrus region of the hippocampus. LFA animals showed significantly lower counts of BrdU+ cells compared to FD+, low folate + succinylsulfathiazole (L5-MTHF+) and HFA animals. HFA animals showed a significant higher amount of BrdU+ cells compared to H5-MTHF+ animals, *indicate significance at *p* = 0.05. **E.** Z-stack photomicrograph of cells co-labeled with the fluorescent neuronal marker NeuN (green) and fluorescent BrdU (red). **F.** Mean and +SEM of the percentage of BrdU+ cells that were also NeuN-positive (NeuN+), and individual data points of **percentage** of BrdU+ cells that were also NeuN+ within the ventral dentate gyrus. L5-MTHF+ animals showed reduced percentage of BrdU+ cells that were also NeuN+ compared to FD+ and L5-MTHF animals. H5-MTHF+ animals showed reduced percentage compared to FD+ and HFA animals. LFA animals also showed reduced percentage compared to L-5MTHF animals, *indicate significance at *p* = 0.05. **G.** Mean and +SEM of the percentage of BrdU+ cells that were also NeuN+, and individual data points of percentage of BrdU+ cells that were also NeuN+ within the dorsal dentate gyrus, no statistically significant differences were found.

L5-MTHF+ animals showed lower percentages of BrdU+ cells co-labelled with NeuN in the ventral region compared to FD+ (*p* = 0.006, Cohen’s *d* = 1.072) and L5-MTHF (*p* < 0.001, Cohen’s *d* = 1.254) animals. L5-MTHF showed a higher percentage compared to LFA animals (*p* = 0.036, Cohen’s *d* = 1.938). H5-MTHF+ animals showed a lower percentage compared to FD+ (*p* = 0.004, Cohen’s *d* = 1.874) and HFA animals (*p* = 0.032, Cohen’s *d* = 1.287). No significant post hoc comparisons were found within the dorsal region, LFA animals however showed a trend towards significance in a higher percentage compared to L5-MTHF+ animals (*p* = 0.084), with a significant interaction between diet and region, F(1, 20) = 2.739, *p* = 0.048, □_p_^2^ = 0.406, but no significant main effect of diet or region (both *p*’s ≥ 0.127). No other pair-wise post hoc comparisons (all *p*’s ≥ 0.100; Figure 4F-G). Chi squared comparisons did not show any differences in percentages of BrdU/NeuN+ cells within treatment groups (Table 3) or between treatment groups (Table 4).

**Table 3.** Description of observed and expected (expected value = (row total x column total)/grand total) % of bromodeoxyuridine (BrdU)-positive (BrdU+) cells that were also NeuN-positive (NeuN+), with individual Chi squared values comparing within diets.

| **Treatment** | **Observed % BrdU**<br>**and NeuN+** |  | **Expected % BrdU and**<br>**NeuN+** |  | **Individual Chi Squared** |  |
| --- | --- | --- | --- | --- | --- | --- |
|  | **Dorsal** | **Ventral** | **Dorsal** | **Ventral** | **Dorsal** | **Ventral** |
| **FD+** | 76 | 86 | 82.69 | 80.21 | 0.45 | 0.46 |
| **L5-MTHF** | 71 | 91 | 82.78 | 80.30 | 1.59 | 1.64 |
| **L5-MTHF+** | 67 | 59 | 64.29 | 62.37 | 0.16 | 0.16 |
| **LFA** | 83 | 77 | 81.59 | 79.15 | 0.04 | 0.04 |
| **H5-MTHF+** | 73 | 72 | 74.02 | 71.81 | 0.00 | 0.00 |
| **HFA** | 83 | 56 | 71.09 | 68.96 | 2.32 | 2.39 |

**Table 4.**
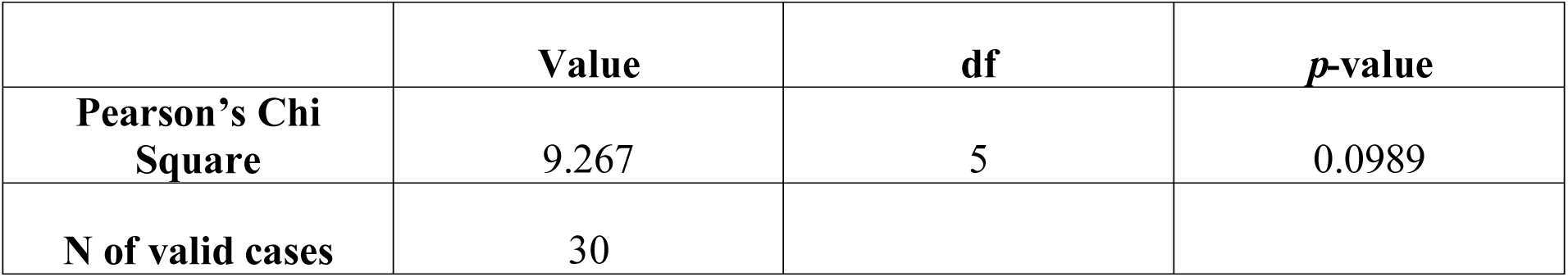
Description of overall Chi square values comparing between diets.

We next calculated a neurogenesis index by multiplying the percentage of BrdU+ cells that also expressed NeuN with the number of BrdU+ cells counted in total as has been done in previous studies^41,56^, and found LFA animals showed a significant reduction in ventral neurogenesis compared to FD+ (p = 0.001, Cohen’s d = 4.412), L5-MTHF (p = 0.004, Cohen’s d = 1.193), and HFA (p < 0.001, Cohen’s d = 2.403) animals. H5-MTHF+ animals showed a significant reduction in ventral neurogenesis compared to FD+ (p < 0.001, Cohen’s d = 2.972) and HFA animals (p < 0.001, Cohen’s d = 2.181). L5-MTHF animals showed a trend towards significance in higher neurogenesis compared to L5-MTHF+ (p = 0.088) within the ventral region. In the dorsal region, HFA animals showed a significant increase in neurogenesis compared to LFA (p < 0.001, Cohen’s d = 1.727) and H5-MTHF+ animals (p < 0.001, Cohen’s d = 1.129), with a significant main effect of diet, F(5, 23) = 2.892, p = 0.036, □_p_^2^ = 0.386 and a main effect of region, F(1, 23) = 25.801, *p* < 0.001, L_p_^2^ = 0.529, but no significant interaction effect (*p* = 0.239). No other pair-wise comparisons were significant (all *p*’s ≥ 0.128; Figure 4A-B).

### 3.5 Ventral DCX-expressing cells positively correlated with relative adrenal mass, plasma homocysteine, and negatively correlated with serum CORT; within low folate without SST and HFA groups, serum CORT negatively correlated with relative adrenal mass

When analyzing all dietary groups together, there was a significant positive correlation between ventral total DCX-expressing cells and relative adrenal mass (mg/100g body mass, r = 0.451, *p* = 0.021; Figure 5A) and plasma homocysteine levels (r = 0.406, *p* = 0.039; Figure 5B), and negatively correlated with serum CORT after dietary treatment (r = −0.463, *p* = 0.017; Figure 5D). No significant correlations emerged when divide by diet groups after Bonferroni corrections (all *p*’s ≥ 0.030). Within the L5-MTHF and HFA group, relative adrenal mass was negatively correlated with serum CORT levels after dietary treatment, r = −0.997, *p* > 0.001, r = −0.995, *p* = 0.005, respectively (Figure 5C). No other significant comparisons were found by diet (*p* ≥ 0.334).

**Figure 5.**
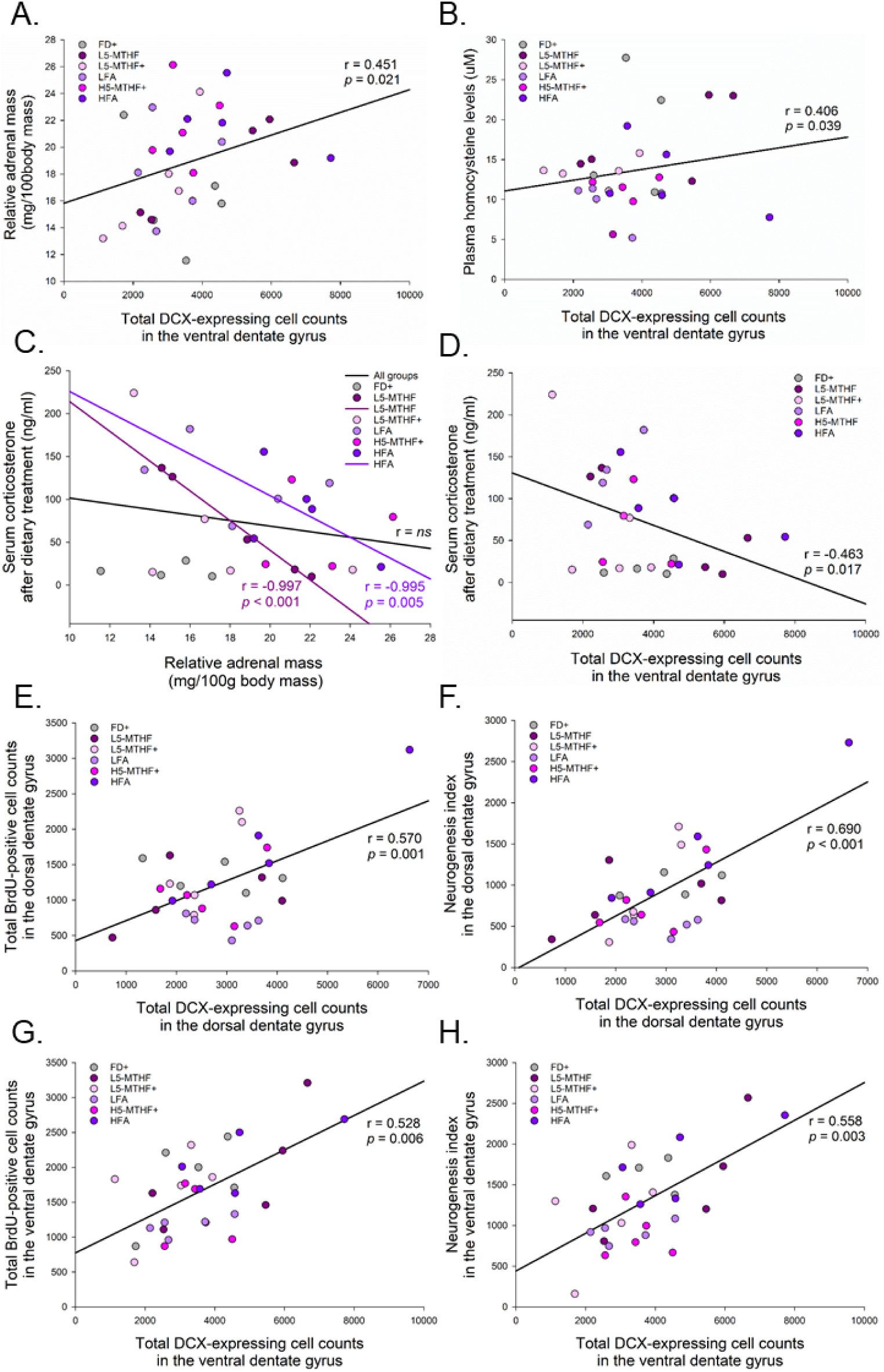
**A.** Correlation between relative adrenal mass and total doublecortin (DCX)-expressing cell counts in the ventral dentate gyrus. A significant positive correlation emerged when comparing all dietary groups together, *p* = 0.05. **B.** Correlation between plasma homocysteine levels and total DCX-expressing cell counts in the ventral dentate gyrus. A significant positive correlation emerged when comparing all dietary groups together, *p* = 0.05. **C.** Correlation between serum corticosterone levels after dietary treatment and relative adrenal mass, where no statistically significant correlation was observed when all dietary animals were group together, *p* = 0.05. When separating animals by dietary interventions, only within the low folate (L5-MTHF) and the high folic acid (HFA) group, did a significant positive correlation emerge after Bonferroni’s correction, *p* = 0.0083. **D.** Correlation between serum corticosterone levels after dietary treatment and total DCX-expressing cell counts in the ventral dentate gyrus. A significant positive correlation emerged when comparing all dietary groups together, *p* = 0.05. **E.** Correlation between bromodeoxyuridine (BrdU)-positive (BrdU+) cells in the dorsal dentate gyrus and total DCX-expressing cell counts in the dorsal dentate gyrus. A significant positive correlation emerged when comparing all dietary groups together, *p* = 0.05. **F.** Correlation between the rates of neurogenesis in the dorsal dentate gyrus and total DCX-expressing cell counts in the dorsal dentate gyrus. A significant positive correlation emerged when comparing all dietary groups together, *p* = 0.05. **G.** Correlation between BrdU+ cells in the ventral dentate gyrus and total DCX-expressing cell counts in the vental dentate gyrus. A significant positive correlation emerged when comparing all dietary groups together, *p* = 0.05. **F.** Correlation between the rates of neurogenesis in the ventral dentate gyrus and total DCX-expressing cell counts in the ventral dentate gyrus. A significant positive correlation emerged when comparing all dietary groups together, *p* = 0.05.

Overall, dorsal total DCX-expressing cells significantly positively correlated with dorsal BrdU+ and dorsal neurogenesis (both *p*’s ≤ 0.001, r = 0.570 and r = 0.690 respectively; Figure 5E-F). Ventral total DCX-expressing cells significantly positively correlated with ventral BrdU+ and ventral neurogenesis (both *p*’s ≤ 0.006, r = 0.528 and r = 0.558 respectively; Figure 5G-H). No significant correlations emerged when comparing BrdU+ cell counts and neurogenesis with serum CORT levels after dietary treatment and relative adrenal mass (all *p*’s ≥ 0.290). Total ventral BrdU+ cell counts and ventral neurogenesis however, showed trends toward positive correlations with plasma homocysteine levels (*p* = 0.089 and *p* = 0.093, respectively; data not shown).

## 4. DISCUSSION

In the present study, we found that high folic acid and low folate (5-MTHF) without the antibacterial SST increased the number of immature neurons (DCX-expressing cells) in the ventral hippocampus compared to the folate deficient diet. In addition, both low folic acid and the high folate (5-MTHF) with SST reduced neurogenesis in the ventral hippocampus. Furthermore, the low 5-MTHF diet group without SST had higher number of DCX-expressing cells in the ventral hippocampus compared to the low 5-MTHF group with SST. The diet-dependent changes in DCX-expressing cells resulted in more mature post-mitotic cells than proliferative cells. However, interestingly the addition of SST, reduced the percentage of BrdU+ cells that also co-expressed NeuN, a marker for mature neurons, suggesting that SST influences cell fate in the ventral dentate gyrus. High folic acid and high 5-MTHF with SST increased adrenal mass compared to folate deficient animals. Low folic acid and high 5-MTHF with SST decreased plasma homocysteine levels compared folate deficient animals. Diet did not influence the percent change in serum CORT levels across treatment but ventral DCX-expressing cells were negatively correlated with CORT levels. Collectively, these data suggest that there are dose-dependent effects of 5-MTHF and folic acid on ventral hippocampal neurogenesis in the adult female rat, and that SST may interfere with these neurogenic effects.

### 4.1 High folic acid and low folate without SST increased the number of immature neurons within the ventral hippocampus; low folic acid and high folate decreased neurogenesis within the ventral hippocampus

Here, we show that high folic acid and low 5MTHF without SST increased the number of immature neurons in the ventral dentate gyrus compared to folate deficient diet in adult female rats. However, low folic acid and both doses of 5-MTHF groups with SST (low 5-MTHF+ and high 5-MTHF+) reduced the percentage of BrdU+ cells that was also NeuN+ (28-day old cells) in the ventral hippocampus. Other reports indicate that folate deficiency after 3.5 months of a folate deficient diet + SST treatment decreased cell proliferation and the number of 18-day old BrdU+ cells in male mice^17^. Our results are partially consistent with their data, as a folate deficient diet in females reduced DCX-expressing cells (expressed for ∼21 days) compared to a high folic acid diet. However, intriguingly, neurogenesis was suppressed in high 5-MTHF with SST and low folic acid groups compared to the folate deficient diet but only in the ventral dentate gyrus. Together our data suggests that high folic acid can upregulate neurogenesis in the short-term (immature neurons), but low folic acid and folate diets with SST reduce neurogenesis (BrdU+ cells) in the longer term. We injected BrdU after 1 day of dietary treatment and perfused animals 28 days later to examine 28-day old neurons (BrdU/NeuN). However, DCX-expressing cells were examined after 29 days of dietary treatment, which given they are expressed for up to 21 days in rats^25^ suggests these immature neurons are younger than the BrdU+ cells and were produced after more days of exposure to the diet. Here, we see more post-mitotic than proliferative DCX-expressing cells in the ventral dentate gyrus, particularly when animals were treated with HFA diet and low 5-MTHF without SST. This suggests that these new neurons are more likely to survive into a more mature state. But, given there were no significant increases in neurogenesis as measured by BrdU/NeuN this also suggests that many of these cells may die prior to reaching a final mature status.

Furthermore, low 5-MTHF without SST, significantly increased the number of ventral immature neurons compared to low 5-MTHF with SST, suggesting that SST interfered with the ability of 5-MTHF to increase the number of immature neurons in low folate dose. In addition, new neurons (BrdU/NeuN+ cells) produced after one day of treatment are less likely to survive under diets with SST and low folic acid diet. Thus, we can infer that the addition of SST reduced neurogenesis in the presence of 5-MTHF diet, which may have been due to its effects on the microbiota or the reduction in folic acid. Additionally, high 5-MTHF with SST reduced neurogenesis within the ventral hippocampus, but this effect however cannot be distinguished between folate effects or SST effects due to the lack of a high 5-MTHF without SST group. Although, for the purpose of this study, we were particularly interested in whether SST could interfere with low 5-MTHF diet, future studies should consider the addition of a folate deficient group without SST to determine the effects of anti-bacterial agent alone in this condition, or adding SST to higher does of 5-MTHF diet to determine if there is a dose-dependent response. Overall, our findings suggest strong effects of the antibacterial SST on early stages of neurogenesis, which is somewhat consistent with Mohle et al.^57^, who found seven weeks of antibiotics reduced four-week old BrdU+ cells and neurogenesis in adult female mice. Future studies should consider a longer treatment period or analyzing the number of immature neurons and mature neurons at different timepoints after dietary treatment.

Overall, our results suggest that the introduction of SST (and as a result suppressing gut folic acid synthesis) can interfere with any potential effects of 5-MTHF to promote neurogenesis in the adult female rat. The current study indicates that the introduction of the antibacterial SST to eliminate gut folic acid synthesis negates the effects of natural 5-MTHF on neurogenesis and should not be adopted in studies wishing to increase neurogenesis. In addition, different doses of folic acid and 5-MTHF can affect different aspects of neurogenesis or different population of cells differently within the ventral hippocampus.

### 4.2 The effects of folic acid and folate on neurogenesis, adrenal glands and CORT: Implications for depression

In the present study, we found low folic acid and high 5-MTHF with SST reduced plasma homocysteine levels. It is important to note that we utilized a folate deficient diet as a comparative control group. The purpose of this study was to compare and contrast the effects of different doses of 5-MTHF and folic acid to a folate deficient group. Furthermore, we saw that high folic acid and high 5-MTHF with SST increased adrenal mass (but not serum CORT), and an overall negative correlation between CORT following 5-MTHF/folic acid manipulations to neurogenesis in the ventral dentate gyrus. Our findings were more significant within the ventral dentate gyrus, and this is intriguing because there is indirect evidence to suggest more mRNA of glucocorticoid receptors within the ventral hippocampus, which can be affected by stress exposure^58^. The ventral hippocampus is also highly involved in regulating the HPA-axis as well, particularly in regulating the release of corticotrophin-releasing hormone from the hypothalamus^59^, this relationship is linked to the neurobiology of depression (reviewed in ^60^).

As mentioned earlier, increased folate and folic acid levels have been associated with antidepressant-like properties. Folic acid can either act as an antidepressant alone without co-current pharmacological agent^61^ or as an adjuvant with traditional antidepressants^62^. This effect may be more efficacious for women^62^, and only effective when dietary interventions are prolonged^63^. In the present study, we found more effects of the diet to regulate neurogenesis in the ventral hippocampus. This is of interest as the ventral hippocampus is more involved in regulating stress and anxiety^45,64^. Given that neurogenesis in the hippocampus is important for mediating some effects of antidepressant efficacy^65^ and regulates HPA-axis negative feedback via neurogenesis in ventral dentate^66^, this may have important implications in treatment of depression in females. Furthermore folate boosts antidepressant efficacy and is linked to depression^61,62^, it is possible that folic acid and folate diets can help mediate antidepressant effects by influencing neurogenesis within the ventral hippocampus directly.

In addition, depression is associated with higher levels of homocysteine^67^ and elevated levels of cortisol^31,68^ it may be that these two measures are related in part via their effects following dietary folate. Other reports show in a population of healthy middle-aged men and women, high cortisol levels were negatively associated with serum folate levels^69^, and positively associated with serum homocysteine levels. Elevated levels homocysteine and low folic acid are associated with depressive symptoms in middle-aged cohorts of both men and women^32,33^. In our study, suggestions of associations between increased adrenal mass, reduced serum CORT with increased immature neurons in the ventral hippocampus. However, we also see a positive correlation between the number of ventral immature neurons and homocysteine levels, and only a trend towards a positive correlation with BrdU+ cells. This suggests that homocysteine metabolism may be involved in regulating different aspects of neurogenesis, and may also underlie the differences seen between DCX-expressing cells and BrdU+ cells due to 5-MTHF/folic acid treatment. Furthermore, while higher serum CORT levels overall were negatively correlated to the number of immature neurons, suggesting higher CORT may suppress neurogenesis, this effect is not however, affected by diet. Still, the ability of folic acid and 5-MTHF to modulate the different aspects of neurogenesis in the ventral hippocampus suggests a potential mechanism by which dietary folate could modify mood even without the modulation of CORT. In the present study, our dietary treatment was for 29 days, it is possible that with more prolonged exposure of the diets we would see stronger associations between all these variables. While in general, elevated homocysteine levels are associated with cell death and DNA damage that can lead to reduced neurogenesis, both in vitro^22–24^, and in vivo^23^. In older male rats and mice, folate supplementation lasting for at least 8-20 weeks is effective at inducing genomic changes^70,71^. In the present study, supplement duration for 29 days may not be sufficient for certain folate or folic acid diets to significantly reduce homocysteine levels and to have significant effects on neurogenesis. Thus, more considerations regarding the length of dietary treatment is needed for future studies. It is also crucial to note that here, we utilized healthy female rats to determine whether folate and folic acid could affect biomarkers of depression such as reduced neurogenesis, high levels of CORT or homocysteine. It is also possible that we would see stronger associations between all these variables in animal models of depression.

Another active area of neurogenesis in rodents is the subventricular zone of the olfactory bulbs. Reduced neurogenesis within the olfactory bulbs has been used as a model of depression using adult male rats^72^, and olfactory dysfunction is seen in patients with major depressive disorder^73^ but to our knowledge, no studies have examined the effects of folate diet to influence neurogenesis in this region. Hyperhomocyteinemia have been found to impair both hippocampal and subventricular zone cell proliferation of neural progenitor cells both *in vitro* and *in vivo*^23^. Intriguingly, 14-day folic acid treatment in a rat model of cerebral ischemia increased neurogenesis in the hippocampus and Notch1 signaling, which is critical for SVZ neurogenesis in male rats^37^. Future studies should investigate the effects of folic acid or folate on olfactory bulb neurogenesis.

### 4.3 Folic acid treatment can have different outcomes than 5-MTHF treatment

In the present study, we show some distinct differences in the effects of folic acid vs 5-MTHF on homocysteine clearance and adult hippocampal neurogenesis. In the literature, there are more studies utilizing folic acid compared to 5-MTHF, likely due to the fact that folic acid is more bioavailable than other forms of folate and more stable^11,74^, the latter trait is beneficial for diet fortification, and thus, more relevant for human studies and animal models. It is important to note that folic acid and other forms of folate such as 5-MTHF act through distinct pathways of metabolism^21^ and that the mode of delivery and amount consumed can change the efficacy of both folic acid and folate^75^. Indeed, in the present study, we show that only animals with low folic acid or high 5-MTHF with SST diets showed significantly reduced levels of plasma homocysteine levels compared to the folate deficient group. Furthermore, only animals fed the high folic acid diet and low 5-MTHF without SST, but not the 5-MTHF with SST diets, showed increased number of immature neurons compared to controls. In addition, while low folic acid and high 5-MTHF with SST did not influence the number of immature neurons present in the ventral hippocampus, these groups did show reduced neurogenesis (BrdU/NeuN), without influencing the maturation of immature DCX-expressing neurons. Additionally, high folic acid increased the number of immature neurons, it did not suppress the number of mature neurons, perhaps indicating that these new immature neurons could have reached maturity if we had continued the treatment for a longer period, or alternatively that it takes a week of dietary treatment to stimulate neurogenesis in this region. Furthermore, in the presence of low doses of 5-MTHF, we found that the antibacterial treatment of SST impeded the ability of 5-MTHF to increase the number of immature neurons. While high 5-MTHF significant reduced neurogenesis, this effect may be confounded by SST treatment. Nevertheless, overall high folic acid was found to be the most effective at increasing the number of immature neurons, particularly those that survived into a more mature state, without suppressing mature neurons than low folic acid and 5-MTHF diets with SST. Again, this suggests that a week of dietary treatment with high folic acid is needed to show stimulation of neurogenesis. Low 5-MTHF diet showed similar effect compared to high folic acid than diets with SST, suggesting that without SST, 5-MTHF diets is somewhat comparable to high folic acid. Thus, collectively these results suggest that folic acid diets have greater efficacy to increase the number of immature neurons and reduce homocysteine levels than 5-MTHF diets, and the addition of SST may be interfering with the effects of 5-MTHF to influence immature neurons. Our data also show that both folic acid and 5-MTHF with SST diets showed different dose-dependent effect on neurogenesis rates and homocysteine clearance.

## 5. CONCLUSIONS

In summary, high folic acid and low 5-MTHF without the antibacterial SST increased the number of immature neurons in neurons that would have been exposed to dietary treatments for a week, but did not significantly affect neurogenesis in neurons produced 24 hours after exposure. Low 5-MTHF without the antibacterial SST also increased the number of immature neurons compared to low 5-MTHF treated with the SST, indicating that SST impaired the ability of 5-MTHF to enhance neurogenesis, perhaps via its effects on the microbiome. Both low folic acid and high 5-MTHF with SST decreased neurogenesis in the ventral dentate gyrus (of cells produced after only 24 hours of exposure to the diet) without influencing the number of immature neurons. High folic acid and low 5-MTHF were the only intervention that increased adrenal mass compared to the folate deficient group but surprisingly higher adrenal mass did not correspond to higher serum CORT levels, which may indicate some disturbances in the HPA axes functioning. Low folic acid significantly reduced plasma homocysteine levels at the end of dietary treatment and significantly reduced neurogenesis. Overall, our findings indicate that natural 5-MTHF and synthetic folic acid may act on different pathways to influence hippocampal neurogenesis and homocysteine metabolism. In the presence of low 5-MTHF, SST blocked any effects on immature neurons. The lack of effects of 5-MTHF diets with SST compared to folate deficient animals suggests that modifications to the gut microbiota (or reductions in folic acid) significantly alter the functionality and efficacy of 5-MTHF to modulate neurogenesis in the hippocampus. The current study suggests that folic acid and 5-MTHF have differential effects on neurogenesis, homocysteine metabolism, adrenal mass and serum CORT release depending on does. Thus, we suggest that folic acid and 5-MTHF are not interchangeable. Furthermore, future research should be cautioned that the use of SST in folate studies may interfere with neuroplasticity.

## ACKNOWLEDGMENTS

The authors would like to thank Lucille Hoover, Nikki Kitay, Robin Richardson, Christine Ausman, and especially Stephanie E. Lieblich for their assistance and contributions throughout the experiment. This work was funded by a Canadian Institutes of Health Research (CIHR) operating grant (MOP142308) to LAMG.

## REFERENCES

1. Scott JM a, Weir DG b. Folic acid, homocysteine and one-carbon metabolism: a review of the essential biochemistry. [Review]. Journal of Cardiovascular Risk. 1998 Aug;5(4):223–7.

2. Blom HJ, Smulders Y. Overview of homocysteine and folate metabolism. With special references to cardiovascular disease and neural tube defects. J Inherit Metab Dis. 2011 Feb;34(1):75–81.

3. D’Anci KE, Rosenberg IH. Folate and brain function in the elderly. Curr Opin Clin Nutr Metab Care. 2004 Nov;7(6):659–64.

4. Rossi M, Amaretti A, Raimondi S. Folate Production by Probiotic Bacteria. Nutrients. 2011 Jan 18;3(1):118–34.

5. Burgoon JM, Selhub J, Nadeau M, Sadler TW. Investigation of the effects of folate deficiency on embryonic development through the establishment of a folate deficient mouse model. Teratology. 2002 May 1;65(5):219–27.

6. Ogbonnaya ES, Clarke G, Shanahan F, Dinan TG, Cryan JF, O’Leary OF. Adult Hippocampal Neurogenesis Is Regulated by the Microbiome. Biological Psychiatry. 2015 Aug 15;78(4):e7–9.

7. De Wals P, Tairou F, Van Allen MI, Uh S-H, Lowry RB, Sibbald B, et al. Reduction in Neural-Tube Defects after Folic Acid Fortification in Canada. New England Journal of Medicine. 2007 Jul 12;357(2):135–42.

8. WHO | Periconceptional folic acid supplementation to prevent neural tube defects [Internet]. WHO. [cited 2019 May 16]. Available from: http://www.who.int/elena/titles/folate_periconceptional/en/

9. Country Profiles-Food Fortification Initiative [Internet]. [cited 2019 May 16]. Available from: http://www.ffinetwork.org/country_profiles/index.php

10. Scaglione F, Panzavolta G. Folate, folic acid and 5-methyltetrahydrofolate are not the same thing. Xenobiotica. 2014 May;44(5):480–8.

11. Sanderson P, McNulty H, Mastroiacovo P, McDowell IFW, Melse-Boonstra A, Finglas PM, et al. Folate bioavailability: UK Food Standards Agency workshop report. British Journal of Nutrition. 2003 Aug;90(02):473.

12. Kelly P, McPartlin J, Goggins M, Weir DG, Scott JM. Unmetabolized folic acid in serum: acute studies in subjects consuming fortified food and supplements. Am J Clin Nutr. 1997 Jun 1;65(6):1790–5.

13. Troen AM, Mitchell B, Sorensen B, Wener MH, Johnston A, Wood B, et al. Unmetabolized folic acid in plasma is associated with reduced natural killer cell cytotoxicity among postmenopausal women. J Nutr. 2006 Jan;136(1):189–94.

14. Mattson MP, Shea TB. Folate and homocysteine metabolism in neural plasticity and neurodegenerative disorders. Trends in Neurosciences. 2003 Mar 1;26(3):137–46.

15. Blom HJ, Shaw GM, Heijer M den, Finnell RH. Neural tube defects and folate: case far from closed. Nat Rev Neurosci. 2006 Sep;7(9):724–31.

16. Kronenberg G, Harms C, Sobol RW, Cardozo-Pelaez F, Linhart H, Winter B, et al. Folate Deficiency Induces Neurodegeneration and Brain Dysfunction in Mice Lacking Uracil DNA Glycosylase. J Neurosci. 2008 Jul 9;28(28):7219–30.

17. Kruman II, Mouton PR, Emokpae RJ, Cutler RG, Mattson MP. Folate deficiency inhibits proliferation of adult hippocampal progenitors. NeuroReport. 2005 Jul 13;16(10):1055.

18. Antony A C. In utero physiology: role of folic acid in nutrient delivery and fetal development. Am J Clin Nutr. 2007 Feb 1;85(2):598S–603S.

19. Cameron HA, McKay RD. Adult neurogenesis produces a large pool of new granule cells in the dentate gyrus. J Comp Neurol. 2001 Jul 9;435(4):406–17.

20. Spalding KL, Bergmann O, Alkass K, Bernard S, Salehpour M, Huttner HB, et al. Dynamics of hippocampal neurogenesis in adult humans. Cell. 2013 Jun 6;153(6):1219–27.

21. Scott J, Rébeillé F, Fletcher J. Folic acid and folates: the feasibility for nutritional enhancement in plant foods. Journal of the Science of Food and Agriculture. 2000 May 15;80(7):795–824.

22. Kruman II, Culmsee C, Chan SL, Kruman Y, Guo Z, Penix L, et al. Homocysteine Elicits a DNA Damage Response in Neurons That Promotes Apoptosis and Hypersensitivity to Excitotoxicity. J Neurosci. 2000 Sep 15;20(18):6920–6.

23. Rabaneda LG, Carrasco M, López-Toledano MA, Murillo-Carretero M, Ruiz FA, Estrada C, et al. Homocysteine inhibits proliferation of neuronal precursors in the mouse adult brain by impairing the basic fibroblast growth factor signaling cascade and reducing extracellular regulated kinase 1/2-dependent cyclin E expression. The FASEB Journal. 2008 Nov;22(11):3823–35.

24. Kruman II, Kumaravel TS, Lohani A, Pedersen WA, Cutler RG, Kruman Y, et al. Folic Acid Deficiency and Homocysteine Impair DNA Repair in Hippocampal Neurons and Sensitize Them to Amyloid Toxicity in Experimental Models of Alzheimer’s Disease. The Journal of Neuroscience. 2002 Mar 1;22(5):1752–62.

25. Brown JP, Couillard□Després S, Cooper□Kuhn CM, Winkler J, Aigner L, Kuhn HG. Transient expression of doublecortin during adult neurogenesis. Journal of Comparative Neurology. 2003 Dec 1;467(1):1–10.

26. Coppen A, Bolander-Gouaille C. Treatment of depression: time to consider folic acid and vitamin B12. J Psychopharmacol. 2005 Jan 1;19(1):59–65.

27. Yan J, Liu Y, Cao L, Zheng Y, Li W, Huang G. Association between Duration of Folic Acid Supplementation during Pregnancy and Risk of Postpartum Depression. Nutrients [Internet]. 2017 Nov 2 [cited 2018 Oct 19];9(11). Available from: https://www.ncbi.nlm.nih.gov/pmc/articles/PMC5707678/

28. Molina-Hernández M, Téllez-Alcántara NP, Olivera-López JI, Jaramillo MT. The folic acid combined with 17-β estradiol produces antidepressant-like actions in ovariectomized rats forced to swim. Progress in Neuro-Psychopharmacology and Biological Psychiatry. 2011 Jan 15;35(1):60–6.

29. Rosa PB, Ribeiro CM, Bettio LEB, Colla A, Lieberknecht V, Moretti M, et al. Folic acid prevents depressive-like behavior induced by chronic corticosterone treatment in mice. Pharmacology Biochemistry and Behavior. 2014 Dec 1;127:1–6.

30. Boldrini M, Hen R, Underwood MD, Rosoklija GB, Dwork AJ, Mann JJ, et al. HIPPOCAMPAL ANGIOGENESIS AND PROGENITOR CELL PROLIFERATION ARE INCREASED WITH ANTIDEPRESSANT USE IN MAJOR DEPRESSION. Biol Psychiatry. 2012 Oct 1;72(7):562–71.

31. Stetler C, Miller GE. Depression and Hypothalamic-pituitary-adrenal Activation: A Quantitative Summary of Four Decades of Research. Psychosomatic Medicine. 2011 Feb 1;73(2):114–26.

32. Sachdev PS, Parslow RA, Lux O, Salonikas C, Wen W, Naidoo D, et al. Relationship of homocysteine, folic acid and vitamin B12 with depression in a middle-aged community sample. Psychological Medicine; Cambridge. 2005 Apr;35(4):529–38.

33. Bottiglieri T, Laundy M, Crellin R, Toone BK, Carney MWP, Reynolds EH. Homocysteine, folate, methylation, and monoamine metabolism in depression. Journal of Neurology, Neurosurgery & Psychiatry. 2000 Aug 1;69(2):228–32.

34. Kim MH, Kim E, Passen EL, Meyer J, Kang S-S. Cortisol and estradiol: Nongenetic factors for hyperhomocyst(e)inemia. Metabolism. 1997 Mar;46(3):247–9.

35. de Souza FG, Rodrigues MDB, Tufik S, Nobrega JN, D’Almeida V. Acute stressor-selective effects on homocysteine metabolism and oxidative stress parameters in female rats. Pharmacology Biochemistry and Behavior. 2006 Oct 1;85(2):400–7.

36. Tagbo IF, Hill DC. Effect of folic acid deficiency on pregnant rats and their offspring. Can J Physiol Pharmacol. 1977 Jun;55(3):427–33.

37. Zhang X, Huang G, Liu H, Chang H, Wilson JX. Folic acid enhances Notch signaling, hippocampal neurogenesis, and cognitive function in a rat model of cerebral ischemia. Nutritional Neuroscience. 2012 Mar 1;15(2):55–61.

38. Tanapat P, Hastings NB, Reeves AJ, Gould E. Estrogen stimulates a transient increase in the number of new neurons in the dentate gyrus of the adult female rat. J Neurosci. 1999 Jul 15;19(14):5792–801.

39. Dominguez-Salas P, Moore SE, Cole D, da Costa K-A, Cox SE, Dyer RA, et al. DNA methylation potential: dietary intake and blood concentrations of one-carbon metabolites and cofactors in rural African women. Am J Clin Nutr. 2013 Jun;97(6):1217–27.

40. Nacher J, Crespo C, McEwen BS. Doublecortin expression in the adult rat telencephalon. Eur J Neurosci. 2001 Aug;14(4):629–44.

41. Snyder JS, Choe JS, Clifford MA, Jeurling SI, Hurley P, Brown A, et al. Adult-Born Hippocampal Neurons Are More Numerous, Faster Maturing, and More Involved in Behavior in Rats than in Mice. J Neurosci. 2009 Nov 18;29(46):14484–95.

42. Plümpe T, Ehninger D, Steiner B, Klempin F, Jessberger S, Brandt M, et al. Variability of doublecortin-associated dendrite maturation in adult hippocampal neurogenesis is independent of the regulation of precursor cell proliferation. BMC Neurosci. 2006 Nov 15;7:77.

43. Taupin P. BrdU immunohistochemistry for studying adult neurogenesis: Paradigms, pitfalls, limitations, and validation. Brain Research Reviews. 2007 Jan 1;53(1):198–214.

44. Wojtowicz JM, Kee N. BrdU assay for neurogenesis in rodents. Nature Protocols. 2006 Aug;1(3):1399–405.

45. Fanselow MS, Dong H-W. Are the dorsal and ventral hippocampus functionally distinct structures? Neuron. 2010 Jan 14;65(1):7–19.

46. Schloesser RJ, Manji HK, Martinowich K. Suppression of Adult Neurogenesis Leads to an Increased HP A Axis Response. Neuroreport. 2009 Apr 22;20(6):553–7.

47. Hassouna I, Ott C, Wüstefeld L, Offen N, Neher RA, Mitkovski M, et al. Revisiting adult neurogenesis and the role of erythropoietin for neuronal and oligodendroglial differentiation in the hippocampus. Molecular Psychiatry. 2016 Dec;21(12):1752–67.

48. Workman JL, Gobinath AR, Kitay NF, Chow C, Brummelte S, Galea LAM. Parity modifies the effects of fluoxetine and corticosterone on behavior, stress reactivity, and hippocampal neurogenesis. Neuropharmacology. 2016 Jun;105:443–53.

49. West MJ, Slomianka L, Gundersen HJ. Unbiased stereological estimation of the total number of neurons in thesubdivisions of the rat hippocampus using the optical fractionator. Anat Rec. 1991 Dec;231(4):482–97.

50. West MJ. New stereological methods for counting neurons. Neurobiol Aging. 1993 Aug;14(4):275–85.

51. Kempermann G, Kuhn HG, Gage FH. More hippocampal neurons in adult mice living in an enriched environment. Nature. 1997 Apr;386(6624):493–5.

52. Eadie BD, Redila VA, Christie BR. Voluntary exercise alters the cytoarchitecture of the adult dentate gyrus by increasing cellular proliferation, dendritic complexity, and spine density. Journal of Comparative Neurology. 2005;486(1):39–47.

53. Tanapat P, Hastings NB, Gould E. Ovarian steroids influence cell proliferation in the dentate gyrus of the adult female rat in a dose-and time-dependent manner. J Comp Neurol. 2005 Jan 17;481(3):252–65.

54. Snyder JS, Choe JS, Clifford MA, Jeurling SI, Hurley P, Brown A, et al. Adult-born hippocampal neurons are more numerous, faster maturing, and more involved in behavior in rats than in mice. J Neurosci. 2009 Nov 18;29(46):14484–95.

55. Leuner B, Caponiti JM, Gould E. Oxytocin stimulates adult neurogenesis even under conditions of stress and elevated glucocorticoids. Hippocampus. 2012 Apr;22(4):861–8.

56. Yagi S, Chow C, Lieblich SE, Galea LAM. Sex and strategy use matters for pattern separation, adult neurogenesis, and immediate early gene expression in the hippocampus. Hippocampus. 2016;26(1):87–101.

57. Möhle L, Mattei D, Heimesaat MM, Bereswill S, Fischer A, Alutis M, et al. Ly6C(hi) Monocytes Provide a Link between Antibiotic-Induced Changes in Gut Microbiota and Adult Hippocampal Neurogenesis. Cell Rep. 2016 31;15(9):1945–56.

58. Romeo RD, Ali FS, Karatsoreos IN, Bellani R, Chhua N, Vernov M, et al. Glucocorticoid Receptor mRNA Expression in the Hippocampal Formation of Male Rats before and after Pubertal Development in Response to Acute or Repeated Stress. NEN. 2008;87(3):160–7.

59. Herman JP, Cullinan WE, Young EA, Akil H, Watson SJ. Selective forebrain fiber tract lesions implicate ventral hippocampal structures in tonic regulation of paraventricular nucleus corticotropin-releasing hormone (CRH) and arginine vasopressin (AVP) mRNA expression. Brain Res. 1992 Oct 2;592(1-2):228–38.

60. Nestler EJ, Barrot M, DiLeone RJ, Eisch AJ, Gold SJ, Monteggia LM. Neurobiology of Depression. Neuron. 2002 Mar 28;34(1):13–25.

61. Brocardo PS, Budni J, Kaster MP, Santos ARS, Rodrigues ALS. Folic acid administration produces an antidepressant-like effect in mice: Evidence for the involvement of the serotonergic and noradrenergic systems. Neuropharmacology. 2008 Feb 1;54(2):464–73.

62. Coppen A, Bailey J. Enhancement of the antidepressant action of fluoxetine by folic acid: a randomised, placebo controlled trial. J Affect Disord. 2000 Nov;60(2):121–30.

63. Almeida OP, Ford AH, Hirani V, Singh V, vanBockxmeer FM, McCaul K, et al. B vitamins to enhance treatment response to antidepressants in middle-aged and older adults: results from the B-VITAGE randomised, double-blind, placebo-controlled trial. British Journal of Psychiatry. 2014 Dec;205(06):450–7.

64. Sahay A, Hen R. Adult hippocampal neurogenesis in depression. Nat Neurosci. 2007 Sep;10(9):1110–5.

65. Santarelli L, Saxe M, Gross C, Surget A, Battaglia F, Dulawa S, et al. Requirement of Hippocampal Neurogenesis for the Behavioral Effects of Antidepressants. Science. 2003 Aug 8;301(5634):805–9.

66. Surget A, Tanti A, Leonardo ED, Laugeray A, Rainer Q, Touma C, et al. Antidepressants recruit new neurons to improve stress response regulation. Molecular Psychiatry. 2011 Dec;16(12):1177–88.

67. Almeida OP, Flicker L, Norman P, Hankey GJ, Vasikaran S, van Bockxmeer FM, et al. Association of Cardiovascular Risk Factors and Disease With Depression in Later Life. The American Journal of Geriatric Psychiatry. 2007 Jun 1;15(6):506–13.

68. Burke HM, Davis MC, Otte C, Mohr DC. Depression and cortisol responses to psychological stress: A meta-analysis. Psychoneuroendocrinology. 2005 Oct;30(9):846–56.

69. Cascalheira JF, Parreira MC, Viegas AN, Faria MC, Domingues FC. Serum Homocysteine: Relationship with Circulating Levels of Cortisol and Ascorbate. Annals of Nutrition and Metabolism. 2008;53(1):67–74.

70. Choi S-W, Friso S, Keyes MK, Mason JB. Folate supplementation increases genomic DNA methylation in the liver of elder rats. British Journal of Nutrition. 2005 Jan;93(1):31–5.

71. Keyes MK, Jang H, Mason JB, Liu Z, Crott JW, Smith DE, et al. Older Age and Dietary Folate Are Determinants of Genomic and p16-Specific DNA Methylation in Mouse Colon. J Nutr. 2007 Jul 1;137(7):1713–7.

72. Yang D, Li Q, Fang L, Cheng K, Zhang R, Zheng P, et al. Reduced neurogenesis and pre-synaptic dysfunction in the olfactory bulb of a rat model of depression. Neuroscience. 2011 Sep 29;192:609–18.

73. Pause BM, Miranda A, Göder R, Aldenhoff JB, Ferstl R. Reduced olfactory performance in patients with major depression. Journal of Psychiatric Research. 2001 Sep 1;35(5):271–7.

74. Ohrvik VE, Witthoft CM. Human Folate Bioavailability. Nutrients. 2011 Apr 18;3(4):475–90.

75. Brouwer IA, van Dusseldorp M, West CE, Steegers-Theunissen RPM. Bioavailability and bioefficacy of folate and folic acid in man. Nutrition Research Reviews. 2001 Dec;14(02):267.

